# A Deep Semi-Supervised Framework for Accurate Modelling of Orphan Sequences

**DOI:** 10.1101/2020.07.13.201459

**Authors:** Lewis Moffat, David T. Jones

## Abstract

Accurate modelling of a single orphan protein sequence in the absence of homology information has remained a challenge for several decades. Although not as performant as their homology-based counterparts, single-sequence bioinformatic methods are not constrained by the requirement of evolutionary information and so have a swathe of applications and uses. By taking a bioinformatics approach to semi-supervised machine learning we develop Profile Augmentation of Single Sequences (PASS), a simple but powerful framework for developing accurate single-sequence methods. To demonstrate the effectiveness of PASS we apply it to the mature field of secondary structure prediction. In doing so we develop S4PRED, the successor to the open-source PSIPRED-Single method, which achieves an unprecedented *Q*_3_ score of 75.3% on the standard CB513 test. PASS provides a blueprint for the development of a new generation of predictive methods, advancing our ability to model individual protein sequences.

## Main

Over the past two decades, sequence-based bioinformatics has made leaps and bounds towards better understanding the intricacies of DNA, RNA, and proteins. Large sequence databases^1^ have facilitated especially powerful modelling techniques that use homology information for a given query sequence to infer aspects of its function and structure^2^. A keen example of this progress is in current methods for protein structure prediction that utilize multiple sequence alignments (MSAs) to accurately infer secondary and tertiary structure^3–5^. Unfortunately, much of this progress has not extended to orphan sequences, a very important but very difficult to model class of sequences which have few to no known homologous sequences^5–7^. Also, even when homologues are available, multiple sequence alignment is often too slow to apply to the entirety of a large sequence data bank, and so improved annotation tools which can work with just a single input sequence are also vital in maintaining resources such as InterPro^8^.

A concurrent development is the recent permeation of deep learning methods into bioinformatics; powerful machine learning models that are extremely data hungry but capable of highly accurate inference^3^. Deep learning approaches have seen success in bioinformatics but progress has been constrained as large labelled biological datasets are not always abundantly available^2^. In many biological settings, acquiring labeled data for even a single example can be very costly, although the data itself is often abundant. A clear example is determining high resolution protein structure data. This is evident in that, at current, there are millions of unannotated sequences in the UniProtKB^1^ but a comparatively much smaller number of structures in the PDB^9^.

Here we present Profile Augmentation of Single Sequences (PASS), a general framework for mapping multiple sequence information to cases where rapid and accurate predictions are required for orphan sequences. This simple but powerful framework draws inspiration from Semi-Supervised Learning (SSL) to enable the creation of massive single-sequence datasets in a way that is biologically intelligent and conceptually simple. SSL methods represent powerful approaches for developing models that utilize both labelled and unlabelled data. Where some recent works^10,11^ have looked to take advantage of unlabelled biological sequence data using unsupervised learning, borrowing from techniques in natural language processing^12, 13^, we instead look to modern SSL methods like FixMatch^14^ for inspiration. These methods have demonstrated that psuedo-labelling, amongst other techniques, can significantly improve model performance^14–16^. Pseudo-labelling techniques use the model being trained to assign artificial labels to unlabelled data, which is then incorporated into further training of the model itself^16^.

PASS uses a bioinformatics-based approach to psuedo-labelling to develop a dataset for a given prediction task before training a predictive single-sequence model. First, a large database of sequences is clustered into MSAs. Each MSA is then used as input to an accurate homology-based predictor. The predictions are then treated as pseudo-labels for a single sequence from the MSA. This allows a large unlabelled set of single sequences to be converted into a training set with biologically plausible labels, that can be combined with real labelled data, for training a deep learning based predictor. As an exemplar of the effectiveness of the PASS framework we apply it to the well explored field of single-sequence secondary structure prediction and achieve unprecedented results in the form of Single-Sequence Secondary Structure PREDictor (S4PRED), the next iteration of PSIPRED-Single, our current method. S4PRED achieves a state-of-the-art *Q3* score of 75.3% on the standard CB513 test set^17^. This performance approaches the first version of the homology-based PSIPRED^18^ and represents a leap in performance for single-sequence based methods in secondary structure prediction (Figure 1).

**Figure 1.**
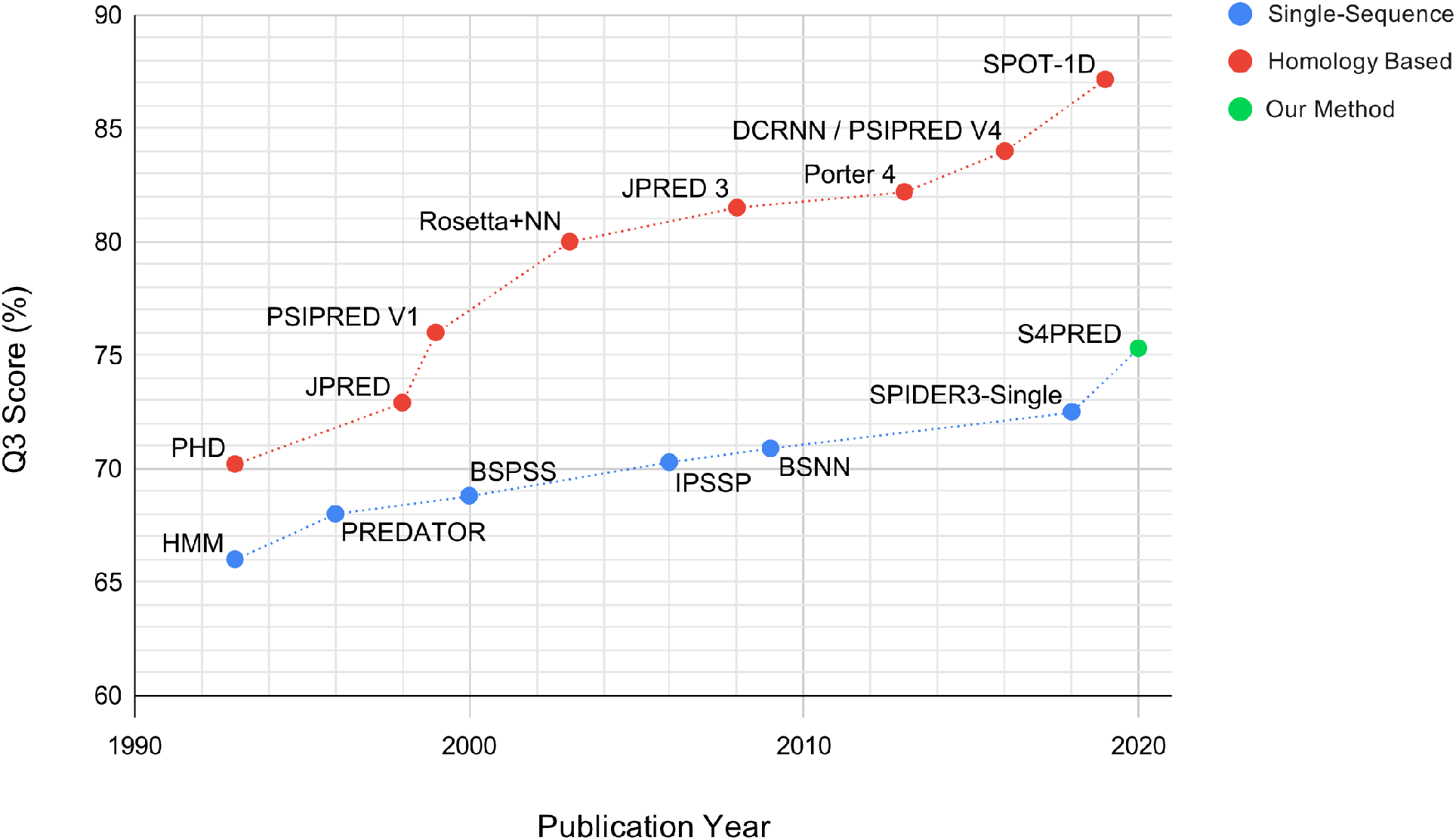
Plot showing reported test *Q*_3_ scores for a range of published secondary structure prediction methods over the previous three decades. This includes single-sequence methods^25,26,30–33^ and homology methods^18,28,34–39^ separately to provide an illustrative view of how single-sequence methods have improved very slowly, compared to homology methods, over time. We include this work, S4PRED, to demonstrate how it is a step upwards in accuracy. In order to avoid conflation with Rosetta *ab initio,* we use the name Rosetta + Neural Network (Rosetta+NN) in this figure to refer to the work of Meiler & Baker^36^.

In the past two decades secondary structure prediction has become an invaluable tool across the cutting edge of protein science, particularly in areas like cryo-electron microscopy^19,20^, tertiary structure prediction^5^, and protein design^21^. Starting from a three class accuracy (Q3) of ~ 76%^18^ in the late 1990’s, our renowned secondary structure prediction tool, PSIPRED, has grown to a current state-of-the-art *Q*_3_ of 84.2%, and is used globally in both experimental and computational research^22^.

PSIPRED, along with other methods, is able to produce high accuracy predictions by leveraging valuable homology information found in MSAs^23^. This is typically done by constructing a MSA for a given query sequence and then converting it into a PSI-BLAST^24^ profile to be used as features for the predictor, along with the original protein sequence^18,23^. This approach is in stark contrast to single-sequence methods, like PSIPRED-Single^22^, that are designed to predict secondary structure based only on a single query sequence, without relying on homology information. Unfortunately, over the past decades, single-sequence methods have been slow to improve relative to homology based methods, as can be seen in Figure 1. Currently, the most performant single-sequence methods achieve low Q3 scores of 71-72%^22,25–27^, where homology based methods are achieving scores of > 84%^22,27,28^ and are approaching a hypothesized theoretical maximum of 88-90%^29^.

Accurate single-sequence prediction enables the modelling of any given sequence without the constraints of homology, which, from both a theoretical and practical perspective, represents an incredibly valuable research prospect with a plethora of use cases. The first and most apparent of these is being able to better model any part of the known protein space, especially given that a quarter of sequenced natural proteins are estimated to have no known homologues^7^ and an even larger portion are inaccessible to homology modelling^5,6,40^. For example, a particularly important area where this is often the case is viral sequence analysis. The structures of viral proteins are often attractive targets for the development of antiviral drugs or the development of vaccines^41^, however, viral sequences tend to be highly diverse and typically have no detectable homologues, making structural modelling difficult^41–43^. Another example is being able to better model the homology-poor “dark proteome”^6^, the contents of which likely holds yet to be discovered functional and structural biology^5^. The value of single-sequence methods also extends outside of natural proteins to areas like *de novo* protein design^21^, where novel sequences and structures typically, by their very design, have no homologues^44^.

Even in the case of a sequence having known homologues, single-sequence methods have many valuable uses. One clear example is in variant effects^43^, where methods like PSIPRED that use MSAs are limited because their predictions for a given sequence will be biased towards a family “average“^2^. Single-sequence methods avoid this bias and have the potential to better model the changes in secondary structure across a family even for highly divergent members. This also extends to being able to better model large single-species insertions that intrinsically have no homology information. Being able to avoid the bias of homology methods could also benefit protein engineering tasks^45^, where the aim may be to generate a sequence that is highly divergent from its homologues.

Not only do single-sequence methods aid in a variety of scientific problems, they also directly tackle research tasks like the protein structure prediction problem. Recent advances in tertiary structure prediction demonstrate highly accurate *ab initio* structure modelling when homologous sequences are available, but successful prediction without homology information remains elusive^4,46^. Single-sequence methods directly address the prediction of protein structure sans homology information, and improved predictors have the potential to lay the groundwork for future steps towards the herculean task of single-sequence tertiary structure prediction.

## Results

### Generating an artificially labelled dataset

For S4PRED, we use the PASS framework to develop a pseudo-labelling approach that is used to generate a large set of single sequences with highly accurate artificial labels. The first step is taking a large set of unlabelled protein sequences clustered as alignments and then removing the clusters containing a small number of sequences. The MSA-based PSIPRED V4^22^ is then used to generate secondary structure predictions for each remaining cluster alignment. The representative sequence for each cluster is used as the target sequence when predicting secondary structure. The target sequence is then kept along with the three-class predictions, and the alignment is discarded. In this way, each cluster produces a single training example, constituting a single sequence and its psuedo-labels.

This approach effectively utilizes a homology-based predictor to provide accurate pseudo-labels for individual unlabelled sequences. PSIPRED generates high accuracy predictions, so it can be inferred that it’s providing highly plausible secondary structure labels. These labels are, therefore, able to provide valuable biological information to the S4PRED model during training. Because each sequence is sampled from a separate cluster, there is also the added benefit of diversity between individual sequences in the dataset. In this work we use the Uniclust30 database^47^ to generate a training set, which, after a rigorous process of benchmarking and cross-validation, contains 1.O8M sequences with pseudo-labels. To accompany the pseudo-labelled sequences, we construct a labelled dataset from protein structures in the PDB^9^. Homology with the test set is evaluated by CATH^48^ classification. The final training and validation sets contain 10143 and 534 sequences respectively.

To train the S4PRED model using both sets of data we adapt the ‘fine-tuning’ approach from recent work of Devlin and collaborators^13^. In the context of S4PRED, fine-tuning consists of first training on the large pseudo-labelled dataset, after which a small amount of additional training is performed with the labelled dataset. Finetuning in this manner provides an effective and regimented training scheme that incorporates both sets of sequences. The S4PRED model itself uses a variant of the powerful AWD-LSTM^49^ model, a recurrent neural network model that uses a variety of regularization techniques.

### The prediction of secondary structure from a single sequence

The final model achieves an average test set Q3 score of 75.3%. This improves the Q3 of PSIPRED-Single by almost 5% (Figure 2A), currently being 70.6%. This is clearly seen in Figure 3A, which shows how the distribution of test set Q3 scores for S4PRED has improved as a whole from PSIPRED-Single scores. In some cases, this has led to a large improvement in prediction accuracy, an example of which is visualized in Figure 3B. Although this represents a significant improvement it is not necessarily a fair comparison as PSIPRED-Single uses a much simpler multi-layer perceptron model^18,22^.

**Figure 2.**
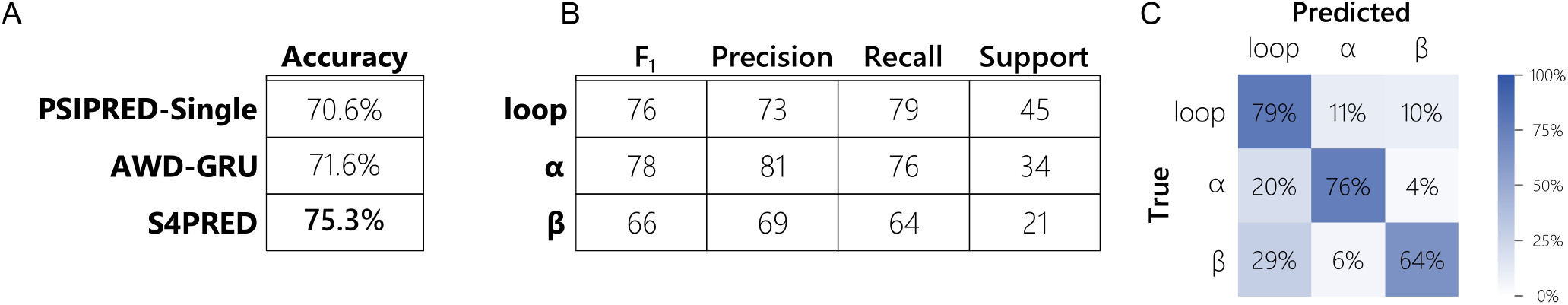
(**A**) Table showing the difference in final accuracy (Q3 score) between the improved S4PRED, the AWD-GRU benchmark, and the current version of PSIPRED-Single on the CB513 test set. (**B**) Table of classification metrics for the S4PRED model test set predictions. These are shown for each of the three predicted class; *α*-helix, *β*-sheet, and loop (or coil). The support is normalized across classes to 100 for clarity – there are a total of 84484 residue predictions in the test set. (**C**) Confusion matrix for the three classes in the S4PRED model test set predictions.

**Figure 3.**
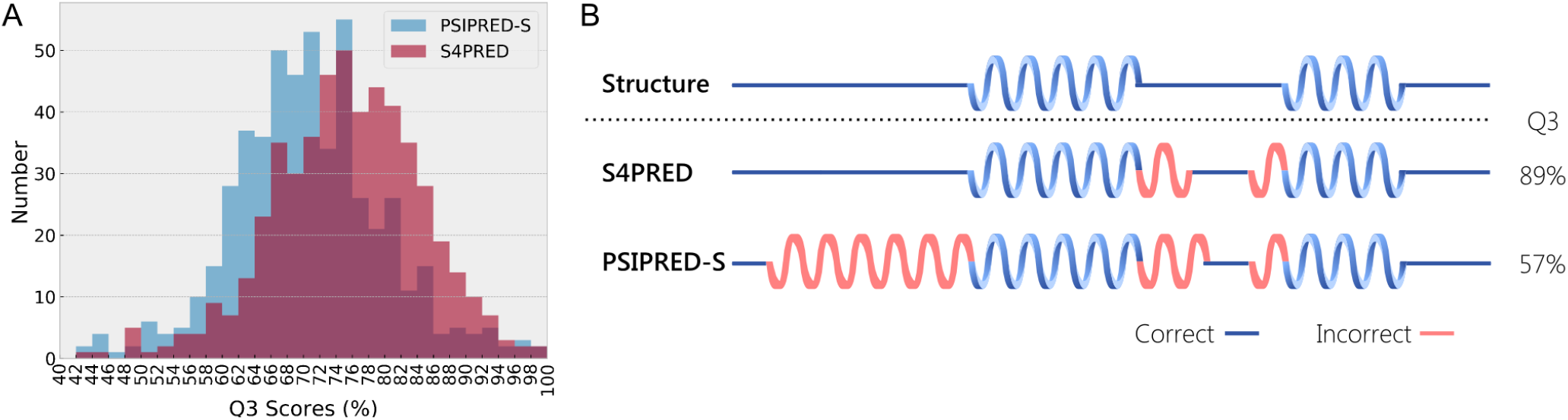
(**A**) Histogram of Q3 scores on the CB513 test set showing the improved results of S4PRED over PSIPRED-Single (PSIPRED-S). (**B**) Example of S4PRED and PSIPRED-Single secondary structure predictions relative to the true structure for the C terminal domain of pyruvate oxidase and decarboxylase (PDB ID 1POW). Although they use the same architecture, S4PRED still exceeds the performance of the AWD-GRU benchmark by a difference in Q3 of almost 4%. Not only is this a large improvement for single-sequence prediction, it directly demonstrates the benefit of the SSL approach.

The most comparable method to date is SPIDER3-Single^26^ which uses a bidirectional LSTM^50^ trained in a supervised manner. This method predicts secondary structure and other sequence information, like solvent accessibility and torsion angles, from a single sequence. SPIDER3-Single uses one model to make preliminary predictions, which are then concatenated with the original input sequence, to be used as input to a second model that produces the final predictions. It reports a *Q*_3_ score of 72.5%, however, this is on a non-standard test set based on a less stringent definition of homology^3^.

To establish an equivalent and informative comparison we provide a second benchmark by training a similar supervised model to SPIDER3-Single which predicts only secondary structure in a standard supervised manner, without a secondary network. This uses the same network architecture as our SSL method but only trains on the labelled sequence dataset. This achieves a Q3 score of 71.6% on CB513. This is a similar result to that achieved in a recent work^27^, which reported a single-sequence *Q*_3_ score of 69.9% and 71.3% on a validation set with a perceptron model and a LSTM-based model respectively. Although the second benchmark used here does not utilize a secondary prediction network like SPIDER3-Single, it is < 1% less performant than SPIDER3-Single’s reported test set performance. Importantly, it provides a direct comparison to S4PRED by using the same model and test set. We use the name AWD-GRU, after the AWD-LSTM variant^49^ used herein, to refer to this benchmark model.

To more precisely determine the benefit that fine-tuning contributes to this performance gain, we tested a model trained on only pseudo-labelled sequences. This achieves a test Q3 score of 74.4%. As is expected, this demonstrates that fine-tuning is a functional approach to combining both datasets that markedly improves prediction by ~ 1%. Aside from the obvious benefit of learning from real labelled data, we speculate that part of the fine-tuning improvement derives from a softening of class decision boundaries. The model trained on only pseudo-labels has a prediction entropy of 0.325, averaged across classes, residues, and sequences. The final model shows a notably higher entropy of 0.548 suggesting that fine-tuning is possibly softening classification probabilities and improving predictions for cases that sit on those boundaries. One clear aspect of S4PRED that should be a focus of future improvement is *β*-strand prediction. Of the three classes it has the lowest *F*_1_ score by a reasonable margin, 0.66 compared to 0.78 and 0.76 for loop and helix respectively (Figure 2B). This is likely due to a combination of being the least represented class in the training set and the most difficult class to predict.

### Data efficiency using the semi-supervised learning approach

Another aspect we wished to investigate was the data efficiency of the SSL approach. We trained the AWD-GRU benchmark model on training sets of different sizes, randomly sampling from the 10143-sequence real-labelled training set. To a good degree, the test set accuracy linearly increases with the logarithm of the real-labelled training set size (*R*^2^ = 0.92), as can be seen in Figure 4. This trend suggests that the SSL approach simulates having trained on a real sequence dataset that is ~x7.6 larger. Under the loose assumption that the ratio of PDB structures to labelled training set size stays the same, there would need to be greater than 1.2M structures in the PDB (as compared to the 162816 entries available as of 04-2020) to achieve the same performance as S4PRED using only real data.

**Figure 4.**
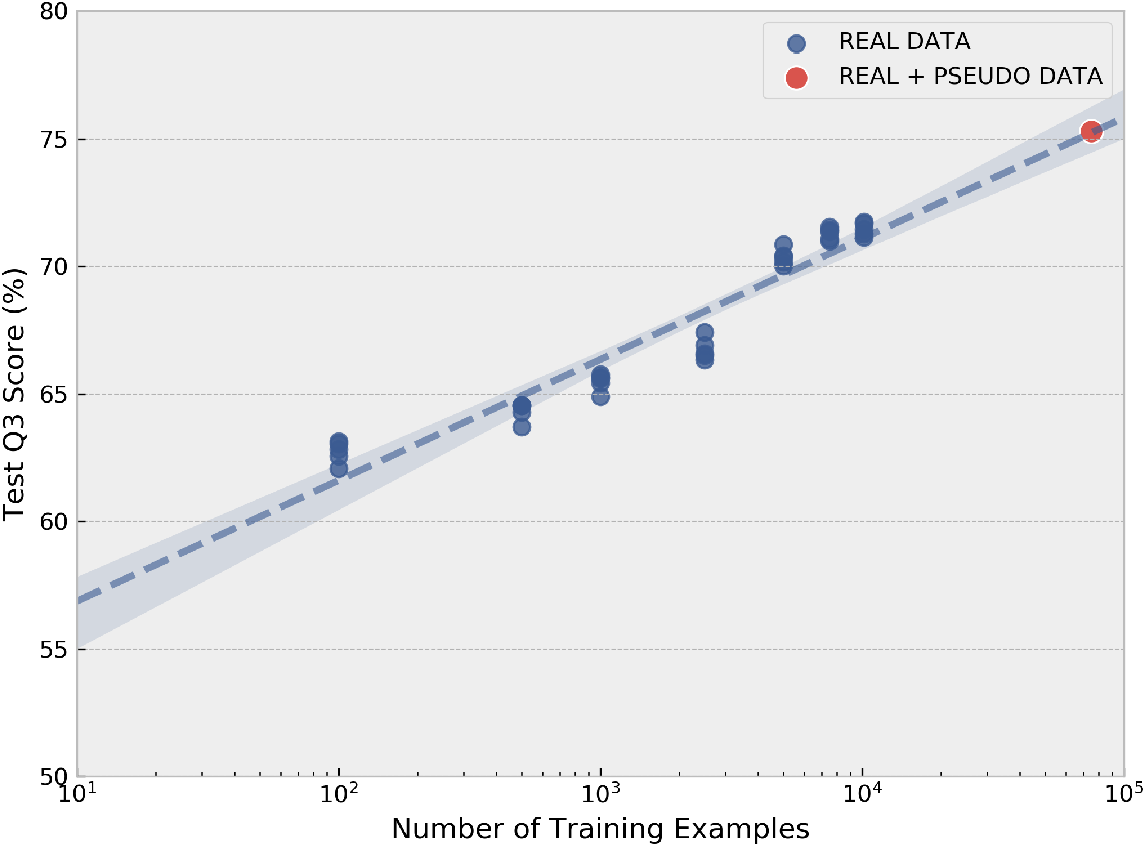
Scatter plot comparing the logarithm of the number of data points compared to trained model accuracy with real labelled sequences. A dashed linear trend line is included. The S4PRED model using real and psuedo-labelled data (75.3%) is included as a single point for comparison.

We also looked to estimate the number of sequences that would be required in UniProt (Swiss-Prot and TrEMBL) and other metagenomic sequence resources^51, 52^ for a PASS-based model to achieve the current performance of the state-of-the-art homology-based PSIPRED. For each single-sequence method in Figure 1, published since the inception of CATH^53^, we find the number of CATH S35 sequence families available the year the method was published. This number servers as a proxy for the number of redundancy-reduced PDB chains that would have been available for generating a dataset. We perform exponential regression between the Q3 scores and the number of CATH S35 sequence families. The S4PRED result is included however 1.08M is used for the number of families. The resulting regression suggests that 25B non-redundant PDBs or sequence clusters would be required for an S4PRED-like model to reach 84%. We then use the average UniClust30 (2016) sequence cluster depth as a multiplicative factor to estimate the number of raw sequences needed. This provides a soft estimate of a minimum of 160 Billion sequences needed for a method based on PASS, like S4PRED, to achieve similar results to current homology based models.

### Single-sequence prediction in context

In this work we consider single-sequence prediction in the strictest sense. This is a model that, for a single example, provides predictions without using information derived from related sequences or evolutionary information. This is an important distinction because using even a small number of homologous sequences improves prediction by several percentage points^33^.

The recently published SPOT-1D^28^ provides a clear example of this phenomenon. Hanson and collaborators^28^ show *Q*_3_ scores of several homology-based models when predicting with low diversity alignments. The criterion for this low diversity is having *N_eff_* < 2 (a measure of alignment diversity, as provided by HHblits^54^). This is reported as *N_eff_* = 1, however, all values are rounded down to the nearest integer. This is clearly not a single-sequence approach. It is also further evidenced in the reported *Q*_3_ scores. Of the methods reported, Porter5^27,55^ achieves the highest *Q*_3_ with 78%, followed by SPOT-1D at 77%. Separate to these results, Porter 5 reports a validation set Q3 of 71.3% when trained on only single sequences without profiles^27^. Ignoring the further potential training set and test set overlap for the values reported in SPOT-1D, this difference in *Q*_3_ clearly demonstrates that using even low diversity alignments is enough to significantly improve predictive performance, over a purely single-sequence approach.

Information from homologous sequences can also improve results by being present in the bias of a trained model. A subtle example of this is in the recent DeepSeqVec model^11^, which trained an unsupervised neural network to produce learned representations of individual sequences from UniRef50^56^. The unsupervised model is subsequently used to generate features which are used to train a second model that predicts secondary structure. This second model achieves a *Q*_3_ score of 76.9% on CB513^11^. Although this two model approach is providing secondary structure predictions for individual sequences, it is not a single-sequence method because the unsupervised model has access to implicit evolutionary information for both the training set and test set sequences. This is partly due to being improperly validated, a split was not performed between the training and test sets. With no split the model is able to learn relationships between test set and training set sequences. It is also due to the training objective of the underlying ELMo language model^57^. The model is able to learn relationships between homologous sequences in a shared latent space, especially given that residue representations are optimized by trying to predict what residue is likely to be found at each position in a given input sequence.

Even if the model uses a small amount of evolutionary information, it still precludes it from being a single-sequence method. The predictions from such a model still benefit from evolutionary information. This not only highlights the difficulty in developing accurate methods that are strictly single-sequence, it also highlights how achieving a *Q*_3_ score of 75.3% with S4PRED represents a step up in performance for single-sequence methods.

## Discussion

Secondary structure prediction from the typical homology-based perspective has improved year-on-year and published *Q*_3_ scores are beginning to rise above 85%. Looking at the history of approaches in the field, the general pattern of methods has remained largely the same; that is, it remains a standard supervised prediction task^23^. In this context, it is easy to assume that the steady rise in model performance seen over the past two decades has resulted from some combination of more powerful classifiers and larger databases. There is a strong argument that a significant majority of the improvements have come from the increase in data available. Model performance is generally a monotonically increasing function of the amount of data and the number of structures in the PDB has increased by an order of magnitude since the turn of the millennium^9^.

It is non-trivial to disentangle the exact relationship between the amount of data available and model performance but the different versions of PSIPRED provide a valuable insight. From an architecture and training perspective, the current version^22^ (V4) remains mostly similar to the original first published model^18^, yet the current version is a state-of-the-art model under strict testing criteria^22^. The primary difference between versions is the much larger available pool of training examples. This suggests strongly that the primary bottleneck on performance has been data availability.

Looking to single-sequence prediction, it stands to reason that methods have improved relatively little over time. Data availability, or more generally the amount of information available to a classifier, appears to be a driving force in performance, and by their very nature single-sequence methods have much less available information. This is likely applicable across many orphan sequence modelling tasks, not just secondary structure predictio^n5,6^. In this work we developed and applied the PASS framework to directly tackle this issue of data availability. This led to the development of S4PRED which, in achieving a leap in single-sequence performance, stands as an exemplar to the effectiveness of the PASS approach. PASS, and S4PRED, leverages a semi-supervised approach to provide a neural network classifier with information from over a million sequences. Not only is this successful, it is also a conceptually simple approach. A homology based method (in this case PSIPRED) is used to generate accurate labels for unlabelled examples. The new example and label pairs are then combined with real-labelled data and used to train a single-sequence based predictor.

S4PRED has achieved significant progress in improving single-sequence secondary structure prediction, but there is still much work to be done. There remains an 8-9% performance gap between S4PRED and current state-of-the-art homology-based methods^23^. Given the importance of data availability, an immediate question that arises is whether the best approach to closing the gap is to simply wait for larger sequence databases to be available in the near future. To an extent, this appears to be a feasible approach. The number of entries in UniProt grows every year^1^ and a massive amount of data is available from clustered metagenomic sequences in databases like the BFD^58,59^.

It is likely that increasing the training set every year will improve performance but to what extent is unknown and the computational cost will correspondingly increase. An increase in training set size will also be dictated by an increase in the number of new families in a database (a sequence cluster being a proxy for a family) and not the number of new sequences. Our estimations suggest that 160 Billion sequences would be required to match homology levels of performance with a PASS method. Given the speed at which sequence databases are growing^1,59^ this is not unreasonable, but unlikely to be within reach in the near future. In short, there is no clear indication that waiting for larger databases will bring single-sequence performance to the level of homology-based prediction, although it will bring some improvement. Instead, a focus on methodological improvements stands to yield the best results.

Looking forward, it is always difficult to speculate what specific methods will result in further improvements. Continuing from the perspective of secondary structure prediction, the field has, in recent years, focused on developing larger and more complex neural networks^23^. There is certainly a benefit to this approach. Prototyping tends to be quick so any improvements found can be shared with the scientific community quickly. For many novel architectures, code is often available and straightforward to adapt into pre-existing secondary structure pipelines due to the pervasive use of auto-differentiation packages like Pytorch^60^ and Tensorflow^61^. A concrete example of this approach would be to adapt multi-headed self-attention to secondary structure prediction and other single sequence prediction fields, having shown significant success in natural language processing^13^.

Unfortunately, there is limited novelty in this overall approach and, most importantly, the results of applying the PASS framework suggest that there are only small gains to be had. Waiting for databases to grow in size, and for the development of more complex network architectures, is unlikely to be the answer. Instead, focusing on developing methods that provide pre-existing models with more prediction-relevant information will likely result in the most significant progress. Admittedly this is an easy concept to pose, and more difficult to execute, but PASS and S4PRED demonstrates that it is possible.

The most obvious approach to this kind of development is to explore further techniques from semi-supervised learning. Methods like data augmentation, that have shown success with image data^14,15^, would be ideal in getting the most out of the data that is available. Unfortunately, it is nontrivial to augment biological sequences even when the structure or function is known which makes data augmentation a difficult approach to pursue^46^. That being said, homologues of a given sequence in the training set can loosely be viewed as biologically valid augmentations of the original target sequence. From this perspective, including multiple pseudo-labelled sequences from each cluster as separate examples, instead of the current method which only includes a single target sequence from each cluster, could be viewed as a proxy for data augmentation. Another approach to improving results may be to train models like S4PRED to predict the class probabilities outputted by the label-providing homology model, instead of predicting the hard class assignments, in a manner similar to Knowledge Distillation^62^. The soft-label information may assist the classifier, although in classification tasks with a small number of classes this information may not contribute significantly. A more general method like MixUp^63^, that is application domain agnostic, might also improve classification by improving the classifiers overall generalizability. Suffice it to say, the semi-supervised approach of PASS brings with it a variety of potential ways to improve performance by directly providing more information to the classifier.

Given the unprecedented success of S4PRED, PASS provides a simple blueprint from which further methods can be developed for modelling orphan sequences. An obvious first step with protein sequences is looking to predict other residue level labels like solvent accessibility^64^ and torsion angle prediction^26^. This could be taken even further and be applied to the nefariously difficult task of protein contact prediction^2^. PASS could also be applied to other biological sequences, such as in the prediction of RNA annotations^65^. Extending PASS to other prediction tasks in the future will also likely be aided by recent efforts to consolidate databases of sequences with pre-calculated predictions of various attributes from a range of tools. One such example being the residue-level predictions provided in DescribePROT^66^. As more of the protein universe is discovered the need for methods that are independent of homology only grows. Methods like S4PRED will hopefully come to represent a growing response to this need, the PASS framework providing a path forward. With this in mind we provide S4PRED as an open source tool and as an option on the PSIPRED web service.

## Methods

### Labelled dataset construction

The first stage in our construction of a labelled dataset is generating a non-redundant set of PDB chains using the PISCES server^67^ with a maximum identity between structures of 70% and a maximum resolution of 2.6Å. This produces a list of 30630 chains, all with a length of 40 residues or more. At the cost of introducing some noise but retaining more examples we do not remove any chains with unlabelled residues.

From this list we then remove any chains that share homology with the test set. We use the standard test set for secondary structure prediction, CB513. Homology is assessed and qualified as having any overlapping CATH^48^ domains at the Superfamily level with any of the sequences in the test set^3^. This removes approximately ⅔ of the chains leaving a total of 10677 from which to generate training and validation sets.

The remaining chains are clustered at 25% identity using `MMseqs2`^68^. From the resulting 6369 clusters, a subset is randomly sampled such that the total sum of their sequences makes up ~ 5% of the 10677 chains. This is to create a validation set that achieves a 95%/5% split between training and validation sets, as well as keeping the validation and test sets similarly sized. This leaves a final split of 10143/534/513 examples for the training, validation, and test sets respectively.

Secondary structures are specified using DSSP^69^. For each residue in each sequence the eight states (H, I, G, E, B, S, T, -) are converted to the standard 3 classes (*Q*_3_) of strand for E & B, helix for H & G, and loop (coil) for the remainder. Protein sequences are represented as a sequence of amino acids, where each residue is represented by one of 21 integers; twenty for the canonical amino acids and one for “X” corresponding to unknown and non-canonical amino acids. Each integer represents an index to an embedding that is learned during the training of the neural network models.

### Pseudo-labelled dataset generation

To assemble a dataset of psuedo-labelled sequences we start with Uniclust30 (Januray 2020 release)^47^. This consists of UniProtKB^1^ sequences clustered to 30% identity, making up 23.8M clusters. Each cluster is then considered as a single potential example for the pseudo-labelled training set. Any cluster can be converted into a target sequence and alignment which can then be passed to PSIPRED to generate high accuracy predictions of secondary structure. These predictions are then one-hot encoded and treated as pseudo-labels with the target sequence providing a single example.

Clusters are filtered from the initial 23.8M Uniclust30 set by removing clusters that are either too short or have too few sequence members. If a cluster has a representative sequence with a length of less than 20 residues or contains less than 10 non-redundant sequences in its alignment it is removed. Applying these restrictions leaves a much smaller set of 1.41M clusters. These are the candidate clusters for generating a training set from which homology with the validation and test sets is to be removed.

### Removal of test set homology from the pseudo-labelled dataset

Models trained on labelled and pseudo-labelled data use the same CB513^17^ test set. This consistency provides a means of directly comparing S4PRED with models trained separately on only labelled data, namely, PSIPRED-Single and the AWD-GRU. The same real-labelled validation set is also used, ensuring that all validation sequences used in this work are structurally non-homologous with the test set.

For the vast majority of clusters, solved structures are not available. This leaves sequence-based approaches to identify and eliminate clusters that share any homology with the test set. It is widely known that using a simple percent identity (e.g. 30%) as a homology threshold between two sequences is inadequate and leads to data leakage^3^. As such we employ a rigorous and multifaceted approach to removing clusters that are homologous to the test set.

The first step is performing HMM-HMM homology searching for each member of CB513 with `HHblits`^54^ using one iteration and an E-value of 10 against the remaining clusters. An accurate means of homology detection, using a high E-value also provides an aggressive sweep to capture any positive matches at the expense of a small number of false hits. One iteration was performed as this was broadly found to return more hits. For the validation set, the same procedure is followed, however the default E-value (1 × 10^-3^) is used with two iterations. All clusters that are matches to the test and validation sets are then removed.

The remaining clusters are copied and combined to create a single large sequence database which is processed with `pFilt`^70^ to mask regions of low amino acid complexity. The test set alignments produced by `HHblits` are used to construct `HMMER`^71^ HMMs which are then used to perform HMM-sequence homology searches against the sequence database using `hmmsearch`. The ‘-max’ flag is used to improve sensitivity and the default e-value is used. All sequences that are positive hits, along with their respective clusters, are removed from the remaining set.

A secondary and overlapping procedure is also performed. Each member of the test set is mapped to one or more Pfam^72^ families by pre-existing annotations. These are found by a combination of SIFTS^73^ and manual searching. From the test set, 17 structures were not found to belong to any Pfam family. For each Pfam family linked to the remaining members of the test set, a list of UniProt sequence IDs is generated. This is extracted from the family’s current UniProt-based Pfam alignment (01-2020) and is used to remove clusters following the same procedure as positive hits from the HMM-sequence search.

In total these methods remove approximately a quarter of the initial 1.41M clusters, leaving a final 1.08M clusters to construct a pseudo-labelled training set. While the fear of data leakage remains ever present, we believe that in the absence of structures this process constitutes a rigorous and exhaustive approach to homology removal.

### Generating pseudo-labels with PSIPRED

A given cluster can provide a sequence with pseudo-labels by first taking its representative sequence as the target sequence and splitting off the remainder of the cluster alignment. This is treated as if it was the target sequence alignment. Both sequence and alignment are then processed using the standard PSIPRED procedure. The three-class secondary structure labels predicted by PSIPRED V4^22^ are then kept along with the target sequence as a single example for the training set. The version of PSIPRED used to generate labels is trained on a set of sequences that are structurally non-homologous with the CB513 test set. This ensures that the pseudo-labels contain no information derived from the test set implicitly through PSIPRED. This procedure is repeated to generate a training set of 1.08M sequences each paired with a sequence of pseudo-labels.

### Model architecture

We use a state-of-the-art recurrent neural network (RNN) from the language modelling domain as a classification model. More specifically we adapt the AWD-LSTM^49^ for secondary structure prediction. The first portion of our model takes a sequence of amino acids encoded as integers and replaces them with corresponding 128-d embeddings that are learned during training and are initialized from 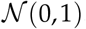. During training a 10% dropout is applied to the embeddings.

The embeddings are fed into a bidirectional gated recurrent unit (GRU)^74^ model with 1024 hidden dimensions in each direction. Here the model differs from the AWD-LSTM which utilizes a long short term memory (LSTM) model with DropConnect^75^ applied to the hidden-to-hidden weight matrices. Our model does the same but utilizes a GRU which we refer to as an AWD-GRU. Unless specified, the weight dropping is set to 50% during training. This model utilizes three layers of AWD-GRUs with 10% dropout applied between each layer during training.

The output of the final recurrent layer is a 2048-d vector at each time step. This is fed into a final linear layer with a log softmax operation to produce the 3-class probabilities at each residue position. These are then used to calculate a negative log likelihood loss using the corresponding one-hot encoded labels. Unlike the original AWD-LSTM we use another popular stochastic gradient descent (SGD) variant, Adam^76^, as an optimizer to minimize the loss and train model parameters.

### S4PRED training with pseudo-labelled data

The first stage in training the S4PRED model is training on the 1.08M pseudo-labelled sequences. For optimization the Adam beta terms are set to *β*_1_,*β*_2_ = {0.9,0.999} with an initial learning rate of 1 × 10^-4^ and a mini-batch size of 256. We also perform gradient clipping with a maximum norm of 0.25. To utilize a batch size of greater than 1 all batches are padded on the fly to the length of the longest sequence in a given batch. The padding symbol has a corresponding embedding and the loss is masked at positions that are padded. Training occurs for up to 10 epochs which typically takes between 48 to 72 hours in total. The performance on the validation set is tested every 100 batches and it is used to perform early stopping.

### Fine-tuning with labelled data

We adapt the methodology presented by Devlin and collaborators^13^ for S4PRED by taking the model trained on pseudo-labelled sequences and performing 1 epoch of training on the 10K labelled sequences. Unlike their method, however, we do not need an additional output layer, having already trained on the semi-supervised secondary structure prediction objective with the psuedo-labelled sequences. For fine-tuning, the batch size is lowered to 32 and the weight drop is set to 0%. All other hyper-parameters are kept the same and the Adam optimizer is reset. The final model is a an ensemble of 5 models fine-tuned with different random seeds, all starting from the same model. Using an ensemble improves prediction by ~ 0.1%.

### Performance benchmarking

Two methods are used to benchmark the results of S4PRED. The first method is the original PSIPRED-Single. Its predictions are generated using the pipeline included with PSIPRED V4. PSIPRED-Single achieves a *Q*_3_ score of 70.6% on CB513. The AWD-GRU model is the second model used for benchmarking. It is trained with the same model architecture and hyper-parameters as S4PRED when it is being trained on the psuedo-labelled set before fine-tuning. However, it only trains on the 10143-sequence set with real labels. This achieves a Q3 score of 71.6% also on CB513.

The data efficiency of the S4PRED method was investigated to estimate the value of training with pseudolabelled data. This was done by training five versions of the AWD-GRU model, each with a different random seed, on different sized subsets of the 10143 real labelled data. Models were trained with 100, 500,1000,2500,5000, 7500, & 10143 examples (a total of 35 models). Each model is tested against CB513 and a linear regression model is fit between the logarithm of the number of points and model Q3 score (*R*^2^ = 0.92). This is visualized in Figure S4. By the linear model, a Q3 score of 75.3% would require 77K real labelled sequences in the dataset.

### Software implementation

All analysis was performed using Python and all neural network models were built and trained using Pytorch^60^. During training, all models used mixed precision which was implemented using the NVIDIA Apex package with the -O2 flag. This was found to improve training speeds with a negligible effect on results. Individual models were trained on a single compute cluster node using an NVIDIA V100 32GB GPU. Upon publication, the S4PRED model and AWD-GRU model with their weights will be released as open source software on the PSIPRED GitHub repository (https://github.com/psipred/) along with documentation. It will also be provided as a part of the PSIPRED web service (http://bioinf.cs.ucl.ac.uk/psipred/).

## Acknowledgements

We thank members of the group for valuable discussions and comments. This work was supported by the European Research Council Advanced Grant ‘ProCovar’ (project ID 695558) and by the Francis Crick Institute which receives its core funding from Cancer Research UK (FC001002), the UK Medical Research Council (FC001002), and the Wellcome Trust (FC001002).

## Author contributions

L.M. and D.T.J. conceived and designed the study and reviewed the manuscript. L.M. carried out the computational work and drafted the manuscript.

## Competing interests

The authors declare no competing interests.

## Notes

### Competing Interest Statement

The authors have declared no competing interest.

### Summary of Updates

Clarified and revised manuscript text.

